# Neural Correlates of Linguistic Collocations During Continuous Speech Perception

**DOI:** 10.1101/2022.03.25.485771

**Authors:** Armine Garibyan, Achim Schilling, Claudia Boehm, Alexandra Zankl, Patrick Krauss

## Abstract

Language is fundamentally predictable, both on a higher schematic level as well as low-level lexical items. Regarding predictability on a lexical level, collocations are frequent co-occurrences of words that are often characterized by high strength of association. So far, psycho-and neurolinguistic studies have mostly employed highly artificial experimental paradigms in the investigation of collocations by focusing on the processing of single words or isolated sentences. In contrast, here we analyze EEG brain responses recorded during stimulation with continuous speech, i.e. audio books. We find that the N400 response to collocations is significantly different from that of non-collocations, whereas the effect varies with respect to cortical region (anterior/ posterior) and laterality (left/right). Our results are in line with studies using continuous speech, and they mostly contradict those using artificial paradigms and stimuli. To the best of our knowledge, this is the first neurolinguistic study on collocations using continuous speech stimulation.

## Introduction

How is natural language processed in the brain and in artificial neural networks? Since decades this issue is tackled from different directions. On the one hand, in experimental neuroscience various neuroimaging techniques are applied to find neural correlates of speech perception in the brain [1]. On the other hand, computational linguistics tries to use computational models to unravel the mystery of language processing [2,3]. While before the 1980s these computational approaches were mainly based on finding and applying strict syntactic rules, the field changed towards statistical natural language processing [2, 4]. Thus, the recent advances in artificial intelligence research were a turning point in the field. Today computational linguistics also called natural language processing (NLP) is based on large text corpora which are used to train artificial neural networks [4]. One field, which made significant progress in the last years and demonstrates the huge impact of this Big Data approach in combination with modern AI systems on natural language processing, is machine translation (MT, [5–7]. How can neuroscience profit from these advances in computational linguistics? Deep artificial neural networks trained on language processing can serve as models for brain function, as argued by Kriegeskorte and Douglas [8, 9], who call that approach “Cognitive Computational Neuroscience” (CCN). In particular, artificial neural networks trained on extensive text corpora can be analyzed to generate hypotheses about important structures and processes involved in language processing. These hypotheses may be tested using neuroimaging data in order to find parallels between artificial and biological neural networks (cf. [10]). Since contemporary AI systems largely lack biological plausibility, existing neural network models have to be made biologically more plausible by, e.g. generating hybrid models from standard machine learning and biologically inspired neuron models [11–14], applying biologically plausible learning rules [15], or biological processing principles such as stochastic resonance and neural noise to make network models more stable [12, 16–19]. This approach is not exclusively useful for neuroscience, but can also be a source of inspiration for novel and more efficient AI approaches [20]. The approach to merge computational linguistics and neurolinguistics is still data limited, which seems counter-intuitive, as modern neuroimaging techniques have a high spatial and temporal resolution, which results in extensive data sets.

However, we have not enough neural data recorded during naturalistic conditions like stimulation with continuous speech to apply the Big Data approach also in neurolinguistics. This is because so far, most neurolinguistic studies have mostly used experimental paradigms that are too simplified, e.g. by focusing on the processing of single words or isolated sentences. As a result, a large number of experimental variables known to affect natural language processing remain very poorly understood. Actually, *“we currently cannot even be sure whether and how benchmark effects from traditional psycho-linguistic studies (e*.*g. word frequency and predictability effects on response times) generalise to more naturalistic situations*.*”* [21]. In contrast, the use of natural language, in particular connected speech, that resembles language as it is used in everyday life offers many advantages over well-controlled, simplified stimuli to study how language is represented and processed in the brain [22–26].

Although for some purposes it might be useful to think of language as a bag of words where the ordering of words does not matter, language is a highly structured system at multiple hierarchical levels where the presence of some linguistic structures can predict or determine the presence of others. Thus, language is fundamentally predictable. For instance, when encountering a ditransitive verb such as *give*, the language user expects the GIVER, the GIVEE and the THING GIVEN, because the argument structure construction implies these participants [27–29]. In addition, for example, when encountering a new object for the first time, one would refer to it using the determiner *the* upon the second encounter because of the definiteness marker.

Another way in which language can be predictable are collocations which are frequently cooccurring word combinations with a high strength of association, e.g.: *go home, annual meeting*, etc. Being an ubiquitous phenomenon, collocations have received much attention from linguistic researchers. There are studies employing both paper-based [30–32] and online/behavioural methods [33–35] of exploring collocations. However, previous studies often looked at collocations in isolation. Among other ways, they would often administer paper-based multiple-choice tasks to reveal participants’ collocational competence or use the phrasal decision task to study the psycholinguistic validity of collocations. Yet, in real life, we do not encounter collocations in isolation. Therefore, this study has attempted to explore collocations using a method that does not rely on physical responses from participants and allows for the presentation of stimuli embedded in sentences and presented naturally. As far as relevant literature is concerned, there is just a handful of neurolinguistic studies of collocations, let alone ERP (event related potentials) studies. In particular, two of these studies are worth mentioning in this respect. Molinaro and Carreiras looked at figurative as well as literal interpretations of Spanish collocations [36]. Using a Rapid Serial Visualization Task (RSVT), they established that collocations in the figurative reading were associated with larger negativities in the N400 time window in comparison with their literal readings suggesting that more processing load is required to integrate the distant meanings in figurative collocations. However, while the title of the paper contains the word ‘collocations’, what the authors mean and explicitly explain in the paper is a broad heterogeneous class of multi-word units, e.g.: collocations, idioms, clichés, proverbs, etc. Thus, the operationalization of collocations in their study is quite different from the strictly linguistic definition of collocations found in the traditional literature on collocations [37–39].

The second study by Hughes [40] comes the closest to our operationalization of collocations in that the difference between collocations and non-collocations is seen as purely quantitative rather than qualitative. So, she uses transitional probability (TP) of 0.01 to distinguish between the two conditions. In a series of experiments, and using the same methodology as Molinaro and Carreiras [36], i.e., RSVT, Hughes [40] claims that non-collocational bigrams are associated with a larger N400 in comparison with collocational bigrams since non-collocations are less expected than collocations, and the effect was right-lateralized. Yet, Hughes [40] has only 15 collocational bigrams which she repeats twice to reach a sufficient number of trials, which is problematic since repetition of the same stimuli can lead to the reduction of the N400 amplitudes [41]. In general, a review of these studies leads to the two following concerns. While it can be argued that RSVT is a more natural task than, for example, a phrasal decision task, the question is whether the task could become even more ecological. Next, the way collocations are operationalized in these studies calls first, for a more linguistic definition of collocation, and second, for a more realistic cut-off point between collocations and non-collocations as far as statistical measures of collocation strength are concerned.

We will conclude these section by presenting our expectations. Thus, given that the N400 is a marker of ease of cognitive processing, with more unpredictable and surprising items showing a larger N400, it was expected that non-collocations will be associated with larger negativities in the N400 time window. As far as the topography is concerned, we did not have any clear expectations because of the mixed findings in the literature. In particular, Hughes [40] reports various distributions in a series of experiments (ranging from anterior through central to posterior scalp distributions). However, as far as lateralization is concerned, we expect that collocations will be processed in the right hemisphere. There are at least two reasons to suggest that. First, in one of the experiments done by Hughes [40], a right-lateralized N400 was reported. Second, given that according to van Lancker and Kempler [42], familiar phrases are processed in the right hemisphere whereas novel ones are processed in the left hemisphere, we hypothesized that collocations (being familiar phrases) will be processed in the right hemisphere, and non-collocations (being novel language) in the left hemisphere.

However in the present study, we find increased N400 amplitudes as a response to collocations in anterior brain regions as well as in more posterior regions in both hemispheres.

## Methods

### Human participants

Participants were 31 (13 females, 18 males) healthy right-handed (augmented laterality index: *μ* = 83.8, *σ* = 20.8) and monolingual native speakers of German aged 20–68 years (*μ* = 27.4 yrs, *σ* = 9.0 yrs). They had normal hearing and did not report any history of neurological illness or drug abuse. They were paid for their participation after signing an informed consent form. Ethical permission for the study was granted by the ethics board of the University Hospital Erlangen (registration no. 161-18 B). For the questionnaire based assessment and analysis of handedness we used the Edinburgh Inventory [43].

### Speech stimuli and natural language text data

As natural language text data, we used the German novel *Gut gegen Nordwind* by Daniel Glattauer (© *Deuticke im Paul Zsolnay Verlag*, Wien 2006) which was published by *Deuticke Verlag*. As speech stimuli, we used the corresponding audio book which was published by *Hörbuch Hamburg*. Both the novel and the audio book are available in stores, and the respective publishers gave us permission to use them for the present and future scientific studies. Book and audio book consist of a total number of 40460 tokens (number of words) and 6117 types (number of unique words). The total duration of the audio book is approximately 4.5 hours. For our study, we only used the first 40 minutes of the audio book, divided into 10 parts of approximately 4 minutes (*μ* = 245 *s, σ* = 39 *s*). This corresponds to approximately 6000 words, or 800 sentences, respectively of spoken language, where each sentence consists on average of 7.5 words and has a mean duration of 3 seconds. In order to avoid cutting the text in the middle of a sentence or even in the middle of a word, we manually cut at paragraph boundaries, which resulted in more meaningful interruptions of the text.

### Stimulation protocol

The continuous speech from the audio book was presented in 10 subsequent parts (cf. above) at a sensory level of approximately 30-60 dB SPL. The actual loudness varied from participant to participant. It was chosen individually to ensure good intelligibility during the entire measurement, but also to prevent it from being unpleasant. Simultaneously with auditory stimulation, a fixation cross at the centre of the screen was presented all the time to minimize artifacts from eye movements. After each audio book part, three multiple-choice questions on the content of the previously presented part were presented on the screen in order to test the participants’ attention. Participants had to answer the questions by pressing previously defined keys on a keyboard. The total duration of the protocol is approximately one hour.

### Generation of trigger pulses with forced alignment

In order to automatically create trigger pulses for both, the synchronization of the speech stream with the EEG recordings, and to mark the boundaries of words for further segmentation of the continuous data streams, *forced alignment* [44–46] was applied to the text and recording. For this study we used the free web service *WebMAUS* [47, 48]. It takes a wave file containing the speech signal, and a corresponding text file as input and gives three files as output: the time tags of word boundaries, a phonetic transcription of the text file, and the time tags of phone boundaries. Even though forced alignment is a fast and reliable method for the automatic phonetic transcription of continuous speech, we carried out random manual inspections in order to ensure that the method actually worked correctly. Although forced alignment is not 100% reliable, manual spot checks found no errors in our alignment. Of course, the high-quality recording of an audio book is among the best possible inputs for such software. For simplicity, we only used the time tags of word boundaries in this study.

### Speech presentation and synchronisation with EEG

The speech signal was presented using a custom made setup. It consists of a stimulation computer connected to an external USB sound device (Asus Xonar MKII, 7.1 channels) providing five analog outputs. The first and second analog outputs are connected to an audio amplifier (AIWA, XA-003), where the first output is connected in parallel to an analog input channel of the EEG data logger in order to enable an exact alignment of the presented stimuli and the recorded EEG signals. In addition, the third analog output of the sound device is used to feed the trigger pulses derived from forced alignment into the EEG recording system via another analog input channel. In doing so, our setup prevents temporal jittering of the presented signal caused by multi-threading of the stimulation PC’s operating system, for instance. The speech sound was presented open field via loudspeakers.

The stimulation software is implemented using the programming language *Python 3*.*6*, together with Python’s sound device library, the *PsychoPy* library [49, 50] for the stimulation protocol, and the *NumPy* library [51] for basic mathematical and numerical operations.

### Electroencephalography and data processing

For EEG recordings we used the actiChamp amplifier from Brain Vision (Brain Products, Brain Vision, Morrisville, USA). The setup has 64 active electrodes, which were recorded with a sampling rate of 2.5 kHz and no further spectral filters, as filtering was performed after the measurement offline, during the evaluation procedure. Electrode impedance was tuned by the application of electrically conductive gel, so that the skin resistance at each electrode location was below 20 kΩ.

Further processing was performed using the Python library *MNE* [52, 53]. The data was bandpassed filtered at 0-10 Hz. Then it was epoched from 200 ms prior stimulus onset to 800 ms post stimulus onset. No baseline correction was applied since in the context of natural speech processing, period before stimulus onset was not period of inactivity. Then artifact rejection based on channel signal amplitude with a threshold of 100*μV* was performed. As a result of artifact rejection, around 39% of trials was lost, which in the context of natural speech processing can be considered as acceptable. Afterwards, evoked data for each participant followed by grand averages across 31 participants were created. The subsequent analysis of brain responses was planned in the following areas: left-anterior area (FC1, FC3, FC5, F1, F3, F5, F7, AF3, AF7), right-anterior area (FC2, FC4, FC6, F2, F4, F6, F8, AF4, AF8), left-posterior (CP1, CP3, CP5, P1, P3, P5, P7) and right-posterior (CP2, CP4, CP6, P2, P4, P6, P8). For this study, we restricted our analyses to sensor space, and did not perform source localization.

### Alignment and segmentation

Since we have both, the original audio book wave file together with the time tags of word boundaries from forced alignment, and the corresponding recordings of two analog auxiliary channels of the EEG, all 64 EEG recording channels could easily be aligned offline with the speech stream. Subsequently, the continuous multi-channel EEG recordings were segmented using the time tags as boundaries.

### Collocations

To find the neural correlates of collocation processing, 13 collocational (e.g., *niedrigem Blutdruck*)^1^ and 13 non-collocational (e.g., *lustlosesten Antwort*) adjective-noun bigrams were selected within the 40-minute audio book fragment. Given that the noun was the critical word to which the brain response was time-locked, and that high-frequency words are associated with reduced N400 effects in comparison with low-frequency words [54], the bigrams were controlled for length and individual word frequency, not phrasal frequency. Operationalization of collcoations is based on a statistical measure of strength of association mutual information (MI). An MI of 5 was taken as a cut-off point between collocations and non-collocations with a buffer zone of approximately 2 units between the conditions.

### Statistical analysis

For statistical analysis, permutation tests were computed for this latency window [55]. The permutation tests were done in *RStudio* [56] using the *Coin* package [57].

## Results

The presentation of collocations compared to non-collocations causes higher N400 negativities in frontal (anterior) brain regions symmetrically in both hemispheres (see Fig. 1, collocations: a1-a3, non-collocations: b1-b3). Additionally, collocations induce a clear negativity in the lateral posterior regions of the left hemisphere (Wernicke’ area), which starts at a latency of 250 ms. However in contrast to that, in the right hemisphere non-collocations cause an increased N400 amplitude compared to collocations indicating that higher level linguistic structures are processed differently in the two hemispheres.

**Figure 1:**
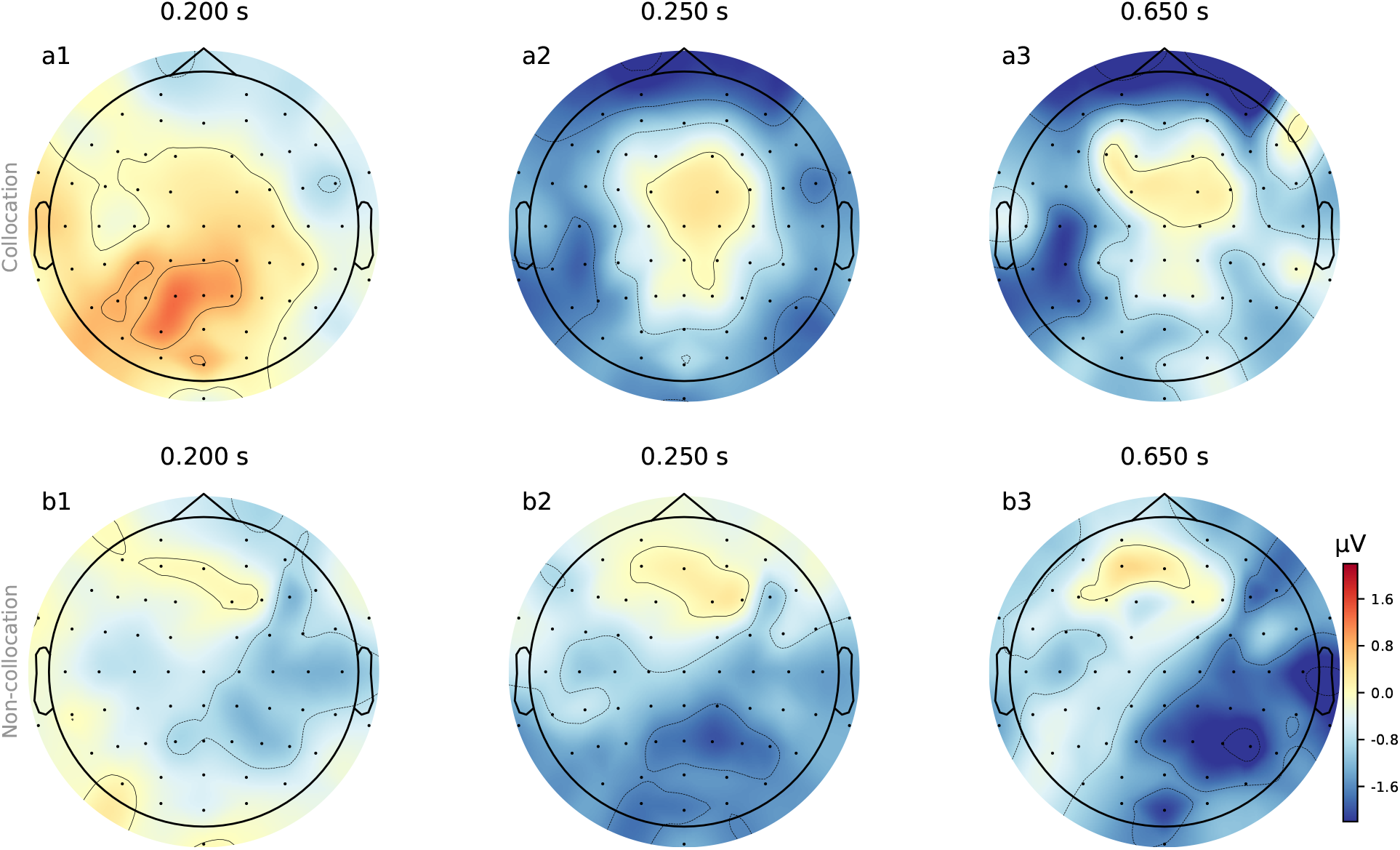
Topomaps of grand averages for collocations and non-collocations. The surface plots show the averaged neural signal (neural activity) of the cerebral cortex averaged over 35 participants for collocations (a1-a3) and non-collocation (b1-b3) stimulation at three different time points (latencies: 200 ms, 250 ms, 650 ms).

In the following, we show the statistical analysis of the negativities within the time-window 300-500 ms after stimulus onset recorded at the four electrode sites (left/right anterior/posterior, for grand averages see Fig. 2, for exemplary subject see 3). The statistical analysis proves that there are indeed larger negativities for collocations in the anterior area (left: -0.467 *μV*, right: -0.644 *μV*) compared to non-collocations (left: -0.093 *μV* ; right: -0.469 *μV*), and larger negativities for non-collocations in the posterior area (left: -0.626 *μV* ; right: -1.198 *μV*) than for collocations (left: -0.567 *μV* ; right:-0.572 *μV*). We restricted our analysis to this time-interval, because we expect the N400 negativity there. This negativity is a marker of unpredictability and surprisal, and therefore a measure of higher processing load.

**Figure 2.**
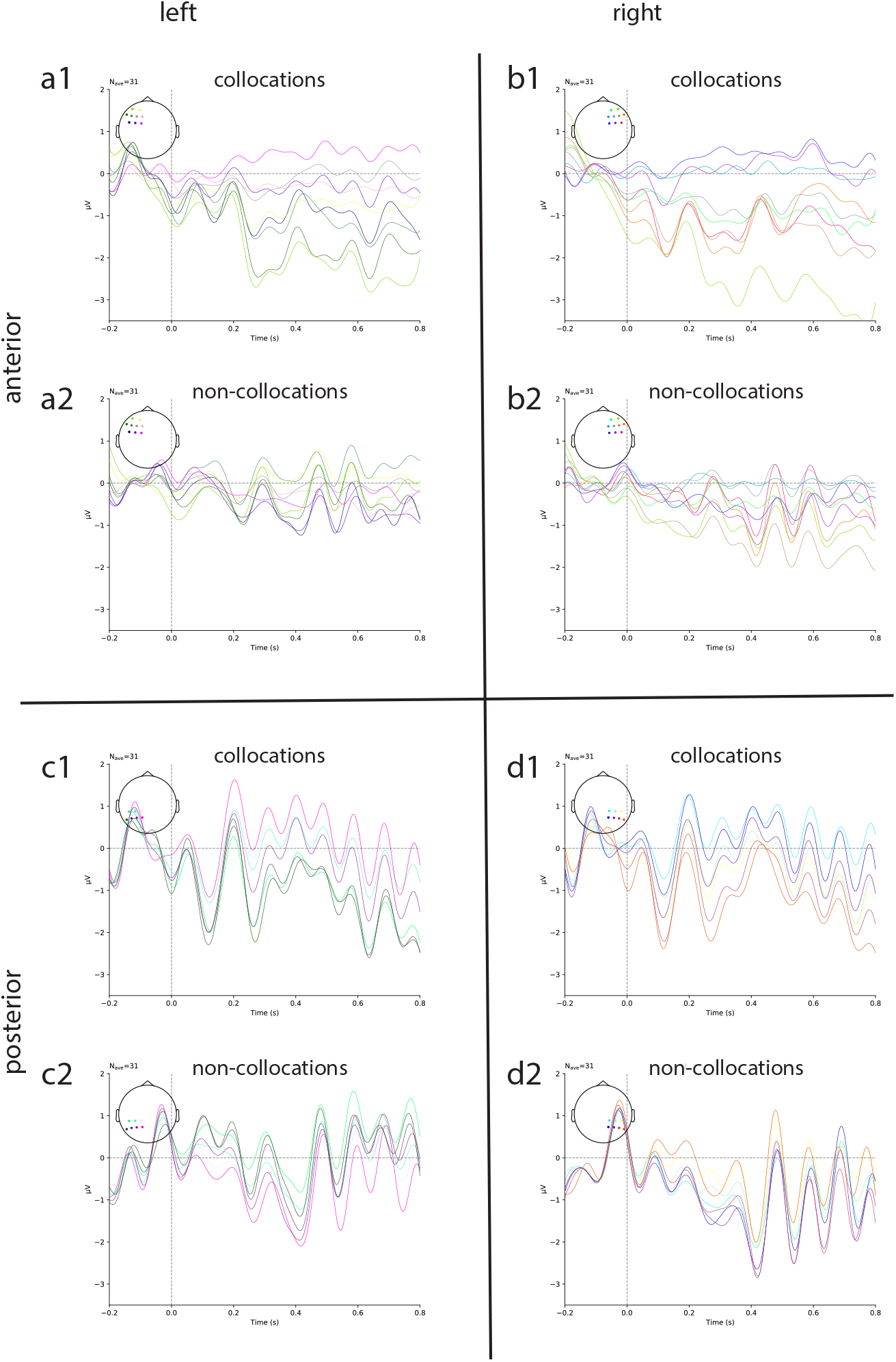
: Grand averages of event related potentials (ERP). a: Averaged neural signal for collocations (a1) and non-collocations (a2) in the left anterior cortex regions; b: Averaged neural signal for the right anterior cortex regions; c-d: Same as a-b for posterior brain regions. (evaluated electrodes are shown as colored markers in the plot inlays, colors of the electrodes correspond to the colors of the plotted curves)

**Figure 3.**
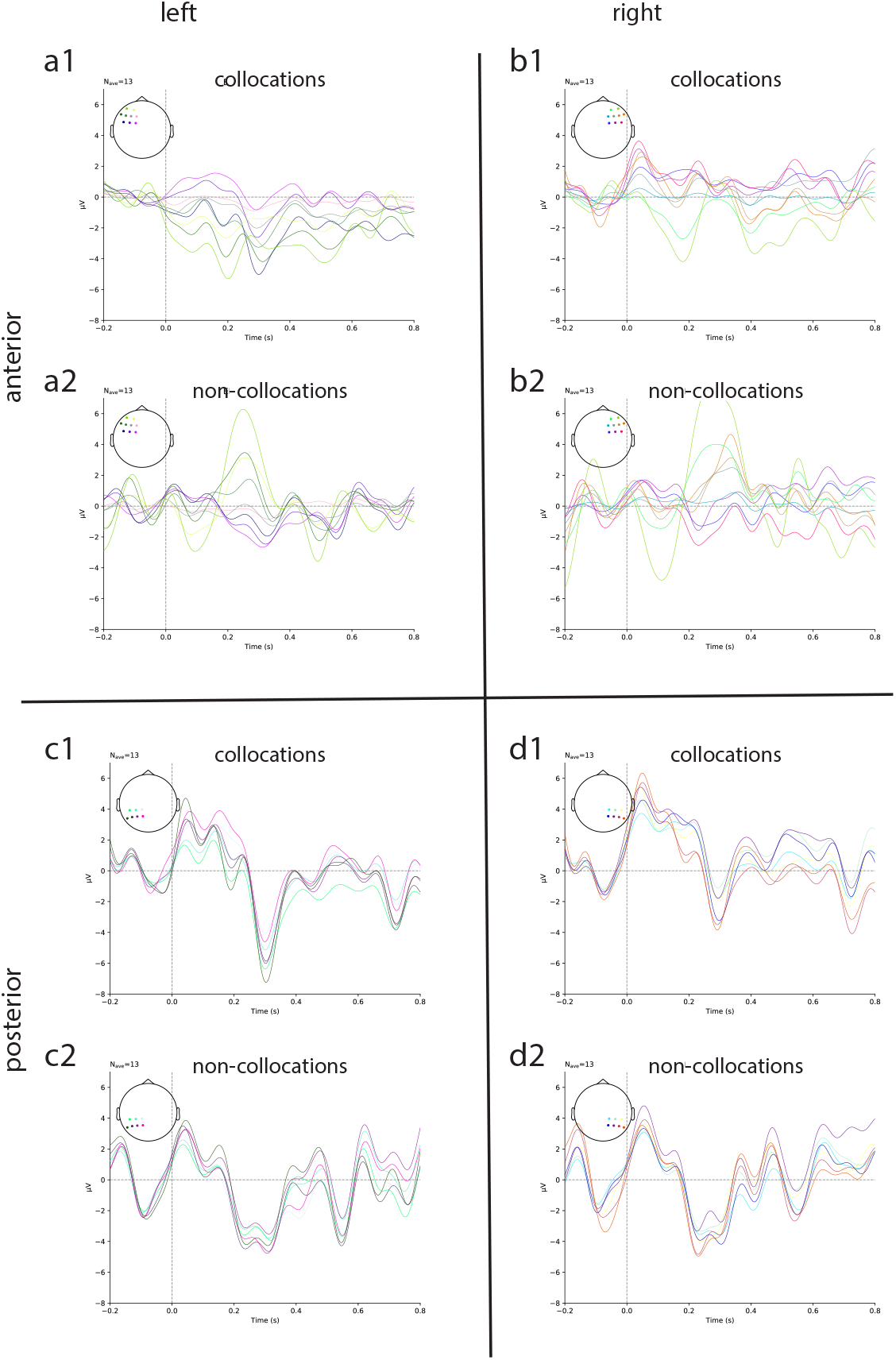
: Event related potentials (ERP) of an exemplary single subject. The plot shows that the effect is not just caused by averaging, but is a real effect also found in single participants. (a: Averaged neural signal for collocations (a1) and non-collocations (a2) in the left anterior cortex regions; b: Averaged neural signal for the right anterior cortex regions; c-d: Same as a-b for posterior brain regions. (evaluated electrodes are shown as colored markers in the plot inlays, colors of the electrodes correspond to the colors of the plotted curves))

According to the results of the permutation tests in the time-interval of 300-500 ms, the observed amplitude differences between collocations and non-collocations were statistically significant at all four electrode sites, i.e., left/right anterior and posterior (*p <* .001 in each case).

Finally, the procedure was repeated to find out whether there was any difference between collocation-processing at the four different recording sites. We show that collocation processing is highly lateralized with the most prominent effect at the anterior regions (*p <* .001).

## Discussion

Given that the N400 is a marker of ease of semantic processing with more unpredictable items showing a (larger) N400, we expected that non-collocations will be associated with larger negativities since they are more unpredictable than collocations. However, in fact, in the N400 time interval, collocations are associated with larger negativities in the anterior area compared to non-collocations (see Figure 1). This finding contradicts the results obtained by Hughes [40] who found a larger N400 wave with anterior scalp distribution for non-collocational bigrams. That our findings are opposite to Hughes [40] who operationalizes collocations in a similar way to ours is controversial. However, the non-collocational bigrams in Hughes [40] are artificially created adjective+noun bigrams, although semantically plausible, that do not appear in the *British National Corpus* (BNC) and were presented in isolated sentences not united by a common context. In the context of the present study, non-collocations appeared in an audio book fragment which is a coherent piece of discourse. Therefore, it would be valid to assume that the non-collocational bigrams in Hughes [40] were more unexpected than the non-collocations in the present study appearing in a natural piece of discourse. However, we also found that collocations were modulated by reduced negative activity in comparison with non-collocations in the N400 time window in the posterior area, which is in line with Hughes [40] who found more negativity for non-collocations in all electrode sites tested, including the posterior area.

Our results can also be supported by Molinaro et al. [36] who studied the effect of unpredictability of complex prepositions in Spanish ending with either a predictable or unpredictable word, e.g.: *in support for/of*. Similarly to us, they found larger negativities (N400-700) to the predictable endings in comparison to the unpredictable ones. Thus, our findings in the anterior area are in line with Molinaro et al. [36] since more predictable units exhibited a larger N400. A word of caution needs to be mentioned, though, due to the fact that unlike the present study, the critical (last) word in Molinaro et al. [36] was a preposition, that is a closed class and a function word. In our study, the critical word was a noun, i.e., an open class and a content word. We are not arguing that this should have necessarily impacted our results in terms of the predictability effects. However, a previous study by Schilling et al. [1] reports a fundamental difference between the processing of content words versus function words in the brain, this fact as well as its possible relation to the interpretation of our results should not be left unnoticed.

Whereas previous literature supporting our findings comes from studies on visual language comprehension, the results from Koskinen and colleagues [58] are especially relevant in the context of the present study since they were also obtained in the context of auditory speech processing. In this work, MEG brain activity was examined during continuous speech processing when participants listened to a 1-hour audio book. The authors found effects of word predictability based on the contextual information in the left hemisphere that mainly involved temporal and frontal brain areas, which overlaps with the left anterior region defined in our study where the second largest difference between conditions (and the largest across the anterior region as well as the left hemisphere) is observed. This finding can be used to argue that collocations as relatively predictable word combinations are predominantly associated with left anterior processing.

Finally, the study that comes the closest to our overall findings is one by Sereno and colleagues [59] who looked at the effects of contextual predictability in reading and found widespread predictability effects in the N400 time window. In addition, the differences were most marked in the left anterior area with more negative amplitudes for high predictability words and more negative-going amplitudes for low predictability words in midline-central and midline-posterior electrode sites. As mentioned earlier, this perfectly matches our results where more predictable items (i.e., collocations) showed more negative-going amplitudes in the left anterior area, and more unpredictable items (i.e., non-collocations) showing negative-going amplitudes in the right-posterior area (see Figure 2).

As far as the laterality of the N400 is concerned, based on the results of the permutation tests, the effect of collocation was significant across all four brain areas. This partially confirms the findings of van Lancker and Kempler [42] who claimed that familiar phrases are processed in the right hemisphere whereas novel language is processed in the left hemisphere because the difference between collocations and non-collocations is the most pronounced in the right-posterior area. However, we also found a statistically significant and large difference between collocations and non-collocations in the left-anterior area, which seems to contradict their findings. We will use our findings to argue that collocations, as defined in the context of the present study, do not share many features with the formulaic language described in van Lancker and Kempler [42] who although do not provide a list of the experimental items, still give a few examples of formulaic language used in the experiment, e.g.: *He’s turning over a new leaf* ; *While the cat is away, the mice will play*. As visible from these items, these are examples of idioms which are both syntactically and semantically fixed, often representing one unit of meaning and therefore having strong imagery. Yet, collocations in our study are regular word combinations that are both syntactically and semantically transparent. That is why it can be argued that the presence of the effect of collocation in the left-anterior area can be explained by the fact that collocations share some features with novel language in that they are analysable multi-word units, whereas the presence of the collocation effect in the right-anterior area can be explained by its idiomaticity.

In addition, it is necessary to point out the importance of modality of the task. Holcomb and Neville [60] studied semantic relatedness in connection with either visual or auditory modality. What they found was larger negativities in the N400 time window in the right hemisphere in the auditory task than in the visual task suggesting that the right hemisphere is responsible for processing prosodic cues in natural speech. Given that the present study is also based on brain responses to natural speech signal, our findings can be argued to be in line with those of Holcomb and Neville [60] because response in both conditions was larger in amplitude in the right hemisphere.

To sum up, our results show that collocations are a psychologically valid phenomenon which was proven by the presence of statistically significant effects in all four electrode sites tested. However, the exact configuration of this effect, e.g., amplitudes, lateralization, remains debatable. We argue that predictability as shown by collocations is modulated by larger negativities in the left anterior area in comparison with non-collocations, but smaller negativities in the right-posterior area. Hence, although we managed to find previous studies that support our findings, it needs to be stated that relating our results to previous literature is challenging because of the many various changes in the configurations of those studies, e.g. item selection criteria, modality of stimuli presentation, task, etc. Kutas and Dale [61] say that ‘N400s do differ in latency and scalp distribution, even within presumably similar experimental tasks’ (p. 222). Thus, small changes of the configurations of a study, leads to different results. Yet, this work has contributed to studies on collocations in many ways. First, to the best of our knowledge, this is the first neurolinguistic study of collocations, let alone in the context of natural speech processing. Also, given that collocations are an ubiquitous phenomenon that we encounter daily in all kinds of discourse, we hope that this study will serve as starting point for more naturalistic studies of collocations, which will lead us closer to the understanding of how these multi-word units are processed in the brain.

However, this study is just a pilot study and further analyses and experiments are needed to gain a more solid data base. Furthermore, the study provides evidence that the results of the neurolinguistics studies have to accompanied by computer simulations, helping to generate hypotheses, which can be tested in the experiments. A lack of these hypotheses makes it nearly impossible to interpret the data. The computational approach can be combined with innovative evaluation techniques based on AI, e.g. dimensionality reduction techniques to account for neural activity spread over the complete cortex [62–64]. This highly interdisciplinary approach based on modern evaluation techniques combined with computer models and strict hypotheses-driven research could potentially solve the problem of low reproducibility between different neurolinguistic and psychological studies [65, 66].

## Acknowledgments

We are grateful to the publishers *Deuticke Verlag* and *Hörbuch Hamburg* for the permission to use the novel and corresponding audio book *Gut gegen Nordwind* by Daniel Glattauer for the present and future studies.

## Author contributions

AG, AS and PK analyzed the data. CB, AZ, AS and PK performed the experiments. AG, AS and PK wrote the manuscript.

## Data availability statement

Data will be made available to other researches on reasonable request.

## Footnotes

1 Grammatical cases left as appearing in the text.

## References

[1] Achim Schilling, Rosario Tomasello, Malte R Henningsen-Schomers, Alexandra Zankl, Kishore Surendra, Martin Haller, Valerie Karl, Peter Uhrig, Andreas Maier, and Patrick Krauss. Analysis of continuous neuronal activity evoked by natural speech with computational corpus linguistics methods. Language, Cognition and Neuroscience, 36(2):167–186, 2021.

[2] Prakash M Nadkarni, Lucila Ohno-Machado, and Wendy W Chapman. Natural language processing: an introduction. Journal of the American Medical Informatics Association, 18(5):544–551, 2011.

[3] G Chowdhury Gobinda. Natural language processing. Annual Review of Information Science and Technology, 37:51–89, 2003.

[4] Dan Klein. The unsupervised learning of natural language structure. Stanford University, 2005.

[5] Lise Volkart, Pierrette Bouillon, and Sabrina Girletti. Statistical vs. neural machine translation: A comparison of mth and deepl at swiss post’s language service. In Proceedings of the 40th Conference Translating and the Computer, pages 145–150, 2018.

[6] Argentina Anna Rescigno, Eva Vanmassenhove, Johanna Monti, and Andy Way. A case study of natural gender phenomena in translation. a comparison of google translate, bing microsoft translator and deepl for english to italian, french and spanish. In CLiC-it, 2020.

[7] Ahmad Yulianto and Rina Supriatnaningsih. Google translate vs. deepl: A quantitative evaluation of close-language pair translation (french to english). AJELP: Asian Journal of English Language and Pedagogy, 9(2):109–127, 2021.

[8] Nikolaus Kriegeskorte and Pamela K Douglas. Cognitive computational neuroscience. Nature neuroscience, 21(9):1148–1160, 2018.

[9] Patrick Krauss and Achim Schilling. Towards a cognitive computational neuroscience of auditory phantom perceptions. arXiv preprint 2010.01914, 2020.

[10] Eric Jonas and Konrad Paul Kording. Could a neuroscientist understand a microprocessor? PLoS computational biology, 13(1):e1005268, 2017.

[11] Richard C Gerum and Achim Schilling. Integration of leaky-integrate-and-fire neurons in standard machine learning architectures to generate hybrid networks: A surrogate gradient approach. Neural Computation, 33(10):2827–2852, 2021.

[12] Achim Schilling, Richard Gerum, Alexandra Zankl, Holger Schulze, Claus Metzner, and Patrick Krauss. Intrinsic noise improves speech recognition in a computational model of the auditory pathway. bioRxiv, 2020.

[13] Paul Stoewer, Christian Schlieker, Achim Schilling, Claus Metzner, Andreas Maier, and Patrick Krauss. Neural network based successor representations of space and language. arXiv preprint 2202.11190, 2022.

[14] Andreas Maier, Harald Köstler, Marco Heisig, Patrick Krauss, and Seung Hee Yang. Known operator learning and hybrid machine learning in medical imaging—a review of the past, the present, and the future. Progress in Biomedical Engineering, 2022.

[15] Richard C Gerum, André Erpenbeck, Patrick Krauss, and Achim Schilling. Sparsity through evolutionary pruning prevents neuronal networks from overfitting. Neural Networks, 2020.

[16] Patrick Krauss, Konstantin Tziridis, Achim Schilling, and Holger Schulze. Cross-modal stochastic resonance as a universal principle to enhance sensory processing. Frontiers in neuroscience, 12:578, 2018.

[17] Patrick Krauss, Konstantin Tziridis, Claus Metzner, Achim Schilling, Ulrich Hoppe, and Holger Schulze. Stochastic resonance controlled upregulation of internal noise after hearing loss as a putative cause of tinnitus-related neuronal hyperactivity. Frontiers in neuroscience, 10:597, 2016.

[18] Achim Schilling, Konstantin Tziridis, Holger Schulze, and Patrick Krauss. The stochastic resonance model of auditory perception: A unified explanation of tinnitus development, zwicker tone illusion, and residual inhibition. bioRxiv, 2020.

[19] Zijin Yang, Achim Schilling, Andreas Maier, and Patrick Krauss. Neural networks with fixed binary random projections improve accuracy in classifying noisy data. In Bildverarbeitung für die Medizin 2021, pages 211–216. Springer, 2021.

[20] Demis Hassabis, Dharshan Kumaran, Christopher Summerfield, and Matthew Botvinick. Neuroscience-inspired artificial intelligence. Neuron, 95(2):245–258, 2017.

[21] Olaf Hauk and Béla Weiss. The neuroscience of natural language processing, 2020.

[22] Nai Ding and Jonathan Z Simon. Neural coding of continuous speech in auditory cortex during monaural and dichotic listening. Journal of neurophysiology, 107(1):78–89, 2012.

[23] Lauren J Silbert, Christopher J Honey, Erez Simony, David Poeppel, and Uri Hasson. Coupled neural systems underlie the production and comprehension of naturalistic narrative speech. Proceedings of the National Academy of Sciences, 111(43):E4687–E4696, 2014.

[24] Michael P Broderick, Andrew J Anderson, Giovanni M Di Liberto, Michael J Crosse, and Edmund C Lalor. Electrophysiological correlates of semantic dissimilarity reflect the comprehension of natural, narrative speech. Current Biology, 28(5):803–809, 2018.

[25] Christian Brodbeck, Alessandro Presacco, and Jonathan Z Simon. Neural source dynamics of brain responses to continuous stimuli: Speech processing from acoustics to comprehension. NeuroImage, 172:162–174, 2018.

[26] Fatma Deniz, Anwar O Nunez-Elizalde, Alexander G Huth, and Jack L Gallant. The representation of semantic information across human cerebral cortex during listening versus reading is invariant to stimulus modality. Journal of Neuroscience, 39(39):7722–7736, 2019.

[27] Adele E. Goldberg. Constructions. University of Chicago Press, Chicago & London, 1995.

[28] Adele E. Goldberg. Construction at work: The nature of generalization in language. Oxford University Press, Oxford, 2006.

[29] Adele E. Goldberg. Explain me this: Creativity, competition, and the partial productivity of constructions. Princeton University Press, Princeton, New Jersey, 2019.

[30] Thomas Herbst. What are collocations: Sandy beaches or false teeth? English Studies, 77(4):379–393, 1996.

[31] Nadja Nesselhauf. Collocations in a learner corpus. John Benjamins, Amsterdam/Philadelphia, 2005.

[32] Ewa Dąbrowska. Words that go together: Measuring individual differences in native speakers’ knowledge of collocations. The Mental Lexicon, 9(3):401–418, 2014.

[33] Brent Wolter and Henrik Gyllstad. Frequency of input and L2 collocational processing. Studies in Second Language Acquisition, 35:451–482, 2013.

[34] Sungmook Choi. Processing and learning of enhanced English collocations: An eye movement study. Language Teaching Research, 21(3), 2017.

[35] Kazuko Matsuno. Processing collocations: Do native speakers and second language learners simultaneously access prefabricated patterns and each single word? Journal of the European Second Language Association, 1(1):61–72, 2017.

[36] Nicola Molinaro and Manuel Carreiras. Electrophysiological evidence of interaction between contextual expectation and semantic integration during the processing of collocations. Biological Psychology, 83:176–190, 2010.

[37] Franz Josef Hausmann. Wortschatzlernen ist Kollokationslernen. Praxis des neusprachlichen Unterrichts, 31:395–406, 1984.

[38] John Sinclair. Corpus, concordance, collocation. Oxford University Press, Oxford, 1991.

[39] John Rupert Firth. Descriptive linguistics and the study of English. In Frank R. Palmer, editor, Selected papers of J.R. Firth 1952-59, pages 96–113. Longmans, London, 1956.

[40] Jennifer Hughes. The psychological validity of collocation: Evidence from event-related brain potentials. Lancaster University (unpublished dissertation), Lancaster, UK, 2018.

[41] Mireille Besson and Marta Kutas. The many facets of repetition: A cued-recall and event-related potential analysis of repeating words in same versus different sentence contexts. Journal of Experimental Psychology: Learning, Memory, and Cognition, 19(5):1115–33, 1993.

[42] Diana Roupas Van Lancker and Daniel Kempler. Comprehension of familiar phrases by left-but not right-hemisphere damaged patients. Brain and Language, 32:265–277, 1987.

[43] Richard C Oldfield. The assessment and analysis of handedness: the edinburgh inventory. Neuropsychologia, 9(1):97–113, 1971.

[44] Pedro J Moreno, Christopher F Joerg, Jean-Manuel Van Thong, and Oren Glickman. A recursive algorithm for the forced alignment of very long audio segments. In ICSLP, volume 98, pages 2711–2714, 1998.

[45] Jiahong Yuan and Mark Liberman. Investigating/l/variation in english through forced alignment. In Tenth Annual Conference of the International Speech Communication Association. Citeseer, 2009.

[46] Athanasios Katsamanis, Matthew Black, Panayiotis G Georgiou, Louis Goldstein, and Shrikanth Narayanan. Sailalign: Robust long speech-text alignment. In Proc. of workshop on new tools and methods for very-large scale phonetics research, 2011.

[47] Florian Schiel. Automatic phonetic transcription of non-prompted speech. 1999.

[48] Thomas Kisler, Uwe Reichel, and Florian Schiel. Multilingual processing of speech via web services. Computer Speech & Language, 45:326–347, 2017.

[49] Jonathan W Peirce. Psychopy — psychophysics software in python. Journal of neuroscience methods, 162(1-2):8–13, 2007.

[50] Jonathan W Peirce. Generating stimuli for neuroscience using psychopy. Frontiers in neuroinformatics, 2:10, 2009.

[51] Charles R Harris, K Jarrod Millman, Stéfan J van der Walt, Ralf Gommers, Pauli Virtanen, David Cournapeau, Eric Wieser, Julian Taylor, Sebastian Berg, Nathaniel J Smith, et al. Array programming with numpy. Nature, 585(7825):357–362, 2020.

[52] Alexandre Gramfort, Martin Luessi, Eric Larson, Denis A Engemann, Daniel Strohmeier, Christian Brodbeck, Roman Goj, Mainak Jas, Teon Brooks, Lauri Parkkonen, et al. Meg and eeg data analysis with mne-python. Frontiers in neuroscience, 7:267, 2013.

[53] Alexandre Gramfort, Martin Luessi, Eric Larson, Denis A Engemann, Daniel Strohmeier, Christian Brodbeck, Lauri Parkkonen, and Matti S Hämäläinen. Mne software for processing meg and eeg data. Neuroimage, 86:446–460, 2014.

[54] Horacio Barber, Marta Vergara, and Manuel Carreiras. Syllable-frequency effects in visual word recognition. Neuroreport, 15(3):545–548, 2004.

[55] Eric Maris and Robert Oostenveld. Nonparametric statistical testing of EEG-and MEG-data. Journal of Neuroscience Methods, 164(1):177–190, 2007.

[56] RStudio Team. RStudio: Integrated development for R. RStudio, Inc., Boston, 2016.

[57] Torsten Hothorn, Kurt Hornik, Mark A. van de Wiel, and Achim Zeileis. Implementing a class of permutation tests: The coin package. Journal of Statistical Software, 28(8):1–23, 2008.

[58] Miika Koskinen, Mikko Kurimo, Joachim Gross, and Aapo Hyvärinen. Brain activity reflects the predictability of word sequences in listenedcontinuous speech. NeuroImage, 219:1–9, 2020.

[59] Sara C. Sereno, Christopher J. Hand, Aisha Shadid, Ian G. Mackenzie, and Hartmut Leuthold. Early EEG correlates of word frequency and contextual predictability in reading. Language, Cognition and Neuroscience, 35(5):625–640, 2020.

[60] Phillip J. Holcomb and Helen J. Neville. Natural speech processing: An analysis using event-related brain potentials. Psychobiology, 19(4):286–300, 1991.

[61] Marta Kutas and Anders Dale. Electrical and magnetic readings of mental functions. In Michael D. Rugg, editor, Cognitive Neuroscience, pages 197–242. Psychology Press, Hove East Sussex, UK, 1997.

[62] Patrick Krauss, Claus Metzner, Achim Schilling, Konstantin Tziridis, Maximilian Traxdorf, Andreas Wollbrink, Stefan Rampp, Christo Pantev, and Holger Schulze. A statistical method for analyzing and comparing spatiotemporal cortical activation patterns. Scientific reports, 8(1):1–9, 2018.

[63] Patrick Krauss, Achim Schilling, Judith Bauer, Konstantin Tziridis, Claus Metzner, Holger Schulze, and Maximilian Traxdorf. Analysis of multichannel eeg patterns during human sleep: a novel approach. Frontiers in human neuroscience, 12:121, 2018.

[64] Patrick Krauss, Claus Metzner, Nidhi Joshi, Holger Schulze, Maximilian Traxdorf, Andreas Maier, and Achim Schilling. Analysis and visualization of sleep stages based on deep neural networks. Neurobiology of sleep and circadian rhythms, 10:100064, 2021.

[65] Leah Maizey and Loukia Tzavella. Barriers and solutions for early career researchers in tackling the reproducibility crisis in cognitive neuroscience. Cortex, 113:357–359, 2019.

[66] Witold M Hensel. Double trouble? the communication dimension of the reproducibility crisis in experimental psychology and neuroscience. European Journal for Philosophy of Science, 10(3):1–22, 2020.

